# TMS-EEG shows mindfulness meditation is associated with an altered excitation/inhibition balance in the dorsolateral prefrontal cortex

**DOI:** 10.1101/2023.10.27.564494

**Authors:** Gregory Humble, Harry Geddes, Oliver Baell, Jake Elijah Payne, Aron T Hill, Sung Wook Chung, Melanie Emonson, Melissa Osborn, Bridget Caldwell, Paul B Fitzgerald, Robin Cash, Neil W Bailey

## Abstract

**Objective:** Mindfulness meditation is associated with functional brain changes in regions subserving higher order cognitive processes such as attention. However, no research to date has causally probed these areas in meditators using combined transcranial magnetic stimulation (TMS) and electroencephalography (EEG). This study aimed to investigate whether cortical reactivity to TMS differs in a community sample of experienced mindfulness meditators when compared to matched controls

**Methods:** TMS was applied to the left and right dorsolateral prefrontal cortices (DLPFC) of 19 controls and 15 meditators while brain responses were measured using EEG. TMS-evoked potentials (P60 and N100) were analysed, and exploratory analyses using the whole EEG scalp field were performed to test whether TMS-evoked global neural response strength or the distribution of neural activity differed between groups.

**Results:** Meditators were found to have statistically larger P60/N100 ratios in response to left and right hemisphere DLPFC stimulation compared to controls (*p*_FDR_ = 0.004, BF_10_ > 39). No differences were observed in P60 or N100 amplitudes when examined in isolation. We also found preliminary evidence for differences in the distribution of neural activity 269-332ms post stimulation.

**Conclusion:** These findings demonstrate differences in cortical reactivity to TMS in meditators. Differences in the distribution of neural activity approximately 300ms following stimulation suggest differences in cortico-subcortical reverberation in meditators that may be indicative of greater inhibitory activity in frontal regions. This research contributes to our current understanding of the neurophysiology of mindfulness and highlights opportunities for further exploration into the mechanisms underpinning the benefits of mindfulness meditation.

---

Mindfulness meditation is a form of mental training that has been found to improve several aspects of general wellbeing (Gu et al., 2015; Spijkerman et al., 2016), including mental health outcomes (Ellen Braun et al., 2019; Godfrin & van Heeringen, 2010). Previous research has indicated that mindfulness meditation is associated with changes in the physiological structure and function of the brain (Fox et al., 2014; Young et al., 2018), including reductions in the age-related decline of grey matter (Pagnoni & Cekic, 2007) and the strengthening of cognitive processes (Chiesa & Malinowski, 2011; Tang et al., 2015), in particular processes related to attention and executive functions (Bailey et al., 2020; Bailey et al., 2023c; Wang et al., 2020). Regions of the brain that regulate attention, emotion, and representation of the self, may be particularly associated with changes due to repeated mindfulness practice (Hölzel et al., 2011; Tang, 2017); Together these three components are believed to work to promote *‘*enhanced self-regulation*’* (Tang, 2017), with increased activity and connectivity between these regions found in meditators (Taren et al., 2017; van Lutterveld et al., 2017).

However, while there is a substantial body of evidence indicating that higher-level cognitive and psychological processes are altered by mindfulness meditation, research has yet to fully characterise the underlying neural mechanisms. Theoretical perspectives suggest that Mindfulness-based Interventions (MBIs) are more likely to yield positive outcomes post-intervention when they are designed to target the underlying intervention mechanisms, rather than targeting only outcomes (Britton, 2019; Britton et al., 2018). As such, gaining a deeper understanding of the neurophysiological mechanisms that lead to brain changes through the practice of meditation might enable evidence-based modifications to MBIs. These modifications could then directly target specific brain regions, networks, and neurophysiological processes implicated in mindfulness practice.

The present study sought to contribute towards a deeper understanding of the structural and functional neural correlates of mindfulness practice. To achieve this, the study combined Transcranial Magnetic Stimulation-Electroencephalography (TMS-EEG) to probe dynamic neuronal behaviour directly. Neuroimaging studies have identified key brain regions associated with enhanced self-regulation in mindfulness meditation. These include the anterior cingulate cortex (ACC), regions of the pre-frontal cortex (PFC), (particularly the dorsolateral prefrontal cortex; DLPFC) and the insula (Ganesan et al., 2022; Tang, 2017). Studies using fMRI have shown differences between controls and experienced mindfulness meditators in networks underlying other cognitive processes. These processes include *attention* (Allen et al., 2012; Brefczynski-Lewis et al., 2007; Brewer et al., 2011; Creswell et al., 2016; Gotink et al., 2016; Hasenkamp & Barsalou, 2012; Jang et al., 2011; Taren et al., 2017; Tomasino & Fabbro, 2016; Xue et al., 2011), *emotion* (Allen et al., 2012; Gotink et al., 2016; Taren et al., 2015; Xue et al., 2011), and *self-regulation* (Allen et al., 2012; Brewer et al., 2011; Gotink et al., 2016; Tomasino & Fabbro, 2016; Xue et al., 2011). Synthesizing this research, one model described by Ganesan et al. (2022), suggests that during focused attention mindfulness meditation, regions responsible for conscious awareness of target interoceptive sensations activate and compete with regions tied to internal thought processes, such as mind wandering. Consequently, the DLPFC and other executive control regions activate to provide top-down regulation, over-riding the processes that produce mind wandering. This action is proposed to diminish the intensity of internal thought processes, allowing focus to be redirected and sustained on the target interoceptive sensation (Ganesan et al., 2022).

The interactions between different brain regions are crucial for brain function, but another fundamental aspect of neurophysiology is the relative balance between cortical excitatory and inhibitory inputs to cortical pyramidal cells (Zhou & Yu, 2018). At pyramidal cells, neurotransmitters and receptors work together within localised dendritic networks to facilitate the changes that lead to structural and functional modifications (Gulyaeva, 2017). Moreover, the balance of excitatory and inhibitory neurotransmission at these neurons plays a key role in creating and maintaining stable oscillatory activity. This activity is not just essential for neuronal signalling, but also underpins higher-order cognitive functions and the brain’s neuroplastic responses to sensory stimuli (Meunier et al., 2017). Therefore, the observed structural and functional changes in meditators might stem from heightened neuroplasticity, potentially arising from shifts in the balance of excitatory and inhibitory inputs to cortical circuits.

Potential improvements in mental health due to mindfulness meditation might be linked to alterations to excitatory/inhibitory (E/I) ratios within prefrontal cortical regions. However, the exact relationship between cortical E/I ratios and mindfulness meditation remains to be explored. TMS-EEG offers a unique window into this dynamic neuronal behaviour. In this method, TMS delivers a targeted magnetic pulse to specific cortical regions of the brain via the scalp. This pulse induces transient electrical current in superficially located neuronal assemblies, leading to the generation of both excitatory and inhibitory postsynaptic potentials in the cortex, known as EPSPs and IPSPs respectively (for a review, see (Farzan et al., 2016). By recording EEG concurrently following the induction of the transient electrical current via TMS, we can observe highly temporally precise information on the cortical response to TMS stimulation. Such data can shed light on the balance and nature of inhibitory and excitatory cortical responses.

The DLPFC is the most thoroughly characterised TMS-EEG stimulation site beyond the primary motor cortex (M1). When EEG activity is measured after stimulating the DLPFC, the result is consistently reproducible TMS-evoked potentials (TEPs). TEPs manifest as complex waveforms with distinct positive peaks and negative troughs at specific post-stimulation latencies. The P60 and N100 are two especially reproducible TEP components that are thought to be associated with glutamatergic excitatory neurotransmission and GABA_B_ergic mediated inhibitory neurotransmission respectively (Cash et al., 2017; Noda et al., 2017; Rogasch et al., 2015; Rogasch & Fitzgerald, 2013). While evidence suggests that glutamatergic and GABAergic neurotransmissions are not the only factors affecting variations in the P60 and N100, a parsimonious interpretation posits that these components are influenced by both inhibitory and excitatory cortical neural activity, and that TEPs generally reflect cortical reactivity. Given the underlying neural basis for these components, several studies have used the P60 to N100 ratio as a marker of cortical E/I balance (Noda et al., 2017; Voineskos et al., 2019). However, previous research has only used TMS-EEG to assess neural activity in meditators by providing stimulation at M1. This research indicated enhanced GABA_B_ergic inhibitory neurotransmission in experienced meditators (Guglietti et al., 2013). This previous research, however, applied stimulation immediately after a 60 minute meditation session which may suggest their findings reflect a state effect rather than a trait effect (Tang et al., 2016). Further, the inhibitory transmission changes at M1 in meditators may reflect alterations in somatosensory processes, such as changes in self-related interoceptive sensory processing (Tang, 2017), rather than attention and executive function related brain regions which are more likely probed by the DLPFC stimulation in the current study.

Considering the provided background, the study’s primary aim was to ascertain if N100 amplitudes differ between experienced mindfulness meditators and non-meditators. A secondary objective was to examine potential differences in the P60 between these two groups. The third aim was to determine if there were noticeable differences in the N100 to P60 ratio when comparing mindfulness meditators with non-meditators. We hypothesised that the N100 TEP component at the DLPFC would show increased amplitudes in experienced mindfulness meditators compared matched controls. We also hypothesised that this would coincide with a compensatory increase the P60 amplitude. Furthermore, we hypothesised (non-directionally), that the E/I balance (operationally defined as the P60/N100 ratio) would differ in mindfulness meditators when compared to matched controls. In addition to these targeted hypotheses, we conducted exploratory analyses on the EEG data to discern if the global neural response strength or the distribution of active neural sources varied between the two groups.

## Methods

### Participants

Experienced mindfulness meditators and healthy controls between the ages of 18 to 65 were recruited to take part in the study. All subjects gave their informed consent prior to participating in any study activities. Participants were recruited through community advertisements, posters, and flyers. Advertisements were posted on social media platforms and flyers were distributed at meditation centres in Victoria, Australia, and within the broader community. Participants responded to the advertisements via phone or email and were screened for inclusion/exclusion criteria. Respondents were excluded from the study if they had received a clinical diagnosis of a mental health disorder, neurological condition or psychiatric illness (either previously or currently occurring), or were currently taking psychoactive meditation and/or illicit drugs, or had a history of concussion or traumatic brain injury in which they had lost consciousness for more than 10 minutes and/or presented to hospital. In addition, anxiety, depression, other psychopathologies and substance use disorders were screened for using the Beck Anxiety Inventory (BAI; (Beck et al., 1988), the Beck Depression Inventory (BDI; (Beck et al., 1996), and the Mini-International Neuropsychiatric Interview (M.I.N.I. version 5.0; (Sheehan et al., 1998), respectively. Participants who scored 20 or above on the BAI (moderate to severe depression range), 16 or above on the BAI (moderate to severe anxiety range) or screened positive to any neuropsychiatric conditions in the M.I.N.I. were excluded from participating.

Meditator respondents were asked to describe their meditation practice. Participants were eligible to participate if their self-described practice adhered to Kabat-Zinn’s definition of mindfulness as “paying attention in a particular way: on purpose, in the present moment, and non-judgementally” (Zinn, 1994). Additional screening questions were asked to ensure the participants’ meditation practice involved focusing on sensations in the body (e.g., paying attention to the breath or body sensations). To address common methodological issues in meditation research of inadequate control for meditation type and level of experience (Van Dam et al., 2018), the current study employed the following criteria for what could be considered an experienced mindfulness meditation practitioner: Meditator participants, at the time of testing, had to be practicing mindfulness meditation (as per Kabat-Zinn’s definition) for two or more hours per week and must have been doing so for a minimum of two years. These criteria have been used in previous research (Bailey et al., 2019; Payne et al., 2020). Uncertainties regarding meditation experience and quality of practice were resolved through direct consultation with the principal investigator (NWB). Healthy controls were included if they had less than 2 cumulative lifetime hours of any form of meditation practice and met all other inclusion criteria. Meditator participants were instructed to not meditate during the testing session. In addition, participants were instructed to avoid caffeine or alcohol for 24 hours prior to administration of the TMS-EEG protocol. At the start of the session participants completed a TMS safety screen (Rossi et al., 2011), and a questionnaire that obtained demographic data (age, years of formal education, gender, handedness, descriptions of their meditation practice including average duration, length and frequency of meditation sessions and current medication use), after which participants completed the BAI, BDI-II, and the Five Facet Mindfulness Questionnaire (FFMQ). This was followed by verbal administration of the M.I.N.I. The current study was reviewed and approved by the Alfred Hospital and Monash University ethics committees, with informed written consent obtained for all participants in accordance with the Declaration of Helsinki.

Of the total 75 participants recruited, TMS-EEG data were not obtained for 36 participants. Of these 36 participants, data were not collected because individuals declined to receive TMS (n = 29), there were concerns regarding TMS eligibility (n = 1), or time restrictions precluded data collection (n = 4). Two further participants were excluded from data analysis due to scoring above the moderate to high anxiety range in the BAI. For a further three participants TMS-EEG data were excluded due to technical issues during recording. Data were excluded for a further single control participant due to reporting meditation experience above the two-hour lifetime maximum limit (which was revealed after the testing session). The final sample of useable TMS-EEG data consisted of 19 controls aged between 20 and 57 years (10 females, *M_age_* = 29.58 years, *SD_age_* 11.48 years), and 15 meditators aged between 22 and 64 years (5 females, *M_age_* = 35.67 years, *SD_age_* 13.81 years).

### Procedure

Each participant took part in a 45 to 60-minute single pulse TMS-EEG data collection block as part of a more extensive session involving EEG recordings and three computerised cognitive tasks. The average total session duration was approximately three-and-a-half hours and involved the following behavioural tasks; i) the Go-Nogo (in preparation), ii) the Oddball (Payne et al., 2020), and iii) the Attentional Blink (Bailey et al., 2023a-a). Following these tasks, the TMS resting motor threshold (RMT) was obtained, then each participant received 105 single pulses of TMS to the left and right DLPFC (with the order counterbalanced between participants).

RMT was determined using electromyography (EMG). Three electrodes were placed surrounding the First Dorsal Interosseous (FDI) muscle on the contralateral (right) hand to the site of stimulation (left M1) and continuous electrical activity of the muscle was observed using Scope V3.7 (ADInstruments, NZ). TMS pulses were delivered to the motor cortex with increasing intensity until a muscle response was observed. RMT was determined as the intensity at which a 1 mV peak-to-peak amplitude motor evoked potential was evoked in the FDI muscle in at least 3 out of 5 trials (Enticott et al., 2012; Rossini et al., 2015).

After RMT was obtained, participants were provided with 105 single pulses of TMS (delivered at a rate of one pulse every four seconds ±10% jitter to control for expectancy effects) to the left- and right-DLPFCs (L-DLPFC, R-DLPFC), respectively. Stimulation intensity was set at 110% of the RMT. Monophasic stimulation was applied using a Magstim 200 stimulator (Magstim Company Ltd., UK) with a figure-of-eight magnetic coil (wing diameter of 70mm). Pulse delivery was automated using Signal version 3.08 software (Cambridge Electronic Design Ltd., UK) connected to a stimulator trigger box. The coil was placed tangentially on the head and positioned with the centre of the coil above frontal EEG electrodes, namely F3 and F4, in accordance with the international 10-20 system for anatomical positioning of TMS stimulation of targeted brain regions. Electrodes F3 and F4 were chosen based on their spatial proximity to the L-DLPFC and R-DLPFC, respectively (Herwig et al., 2003). The TMS coil handle was pointed towards the rear of the participant and 45 degrees laterally from the mid-sagittal line (in accordance with current conventions; (Enticott et al., 2012; Rossini et al., 2015). To control for auditory evoked potentials due to loud clicking noises associated with delivery of each pulse, white noise was delivered using Neuroscan 10Ω ¼ stereo intra-auricular earphone inserts (Compumedics, Melbourne, Australia). The amplitude of white noise was increased gradually to a level that was loud enough to mask the click noises but still comfortable for the participant. Prior to administration of the TMS protocol participants were asked to remain awake and to keep head movements to a minimum.

Cortical electrical activity was obtained using a 64 channel Compumedics Neuroscan Quik-Cap (Compumedics Ltd., Australia) with Ag/AgCl sintered electrodes in the standard 10-20 spatial format. The online reference electrode was located along the midline between CPz and Cz. Additional electrooculography recordings were obtained using a single supraorbital electrode positioned above the left eye in line with the pupil. ECI-Electrogel (Electro-Cap International, Inc., USA) was applied to each electrode and impedances were kept below 5kΩ throughout the duration of the session. EEG data were recorded using DC-coupled 1000x amplification of the EEG signal, acquired using a SynAmps2 amplifier (SynAmps2, EDIT Compumedics Neuroscan, Texas, USA) and recorded using Neuroscan Acquire software V4.5 (Compumedics Ltd., Australia). As per standard TMS-EEG recording guidelines, EEG was recorded using a high sampling rate of 10kHz (*for review see:* (Farzan et al., 2016). Amplifier saturation during TMS delivery was mitigated by using an operating range of ± 200mV with a DC-2kHz low pass filter applied.

#### EEG data processing

EEG data were processed and analysed offline in Matlab R2018b (The MathWorks, USA) utilising the following toolboxes; EEGLAB (Delorme & Makeig, 2004), FieldTrip (Oostenveld et al., 2011), TESA (Rogasch et al., 2017) and Randomization Graphical User Interface (RAGU; (Koenig et al., 2011). Prior to EEG data analysis, channels dedicated to muscle and eye artifact were removed. Initial processing involved removal of the large amplitude TMS pulse (-5 to 15msec), linear interpolation, and the generation and concatenation of epochs (-1 to 2 seconds around the TMS pulse). Data were downsampled to 1000Hz and the removal of DC offset was achieved by baseline correcting each epoch to -500 to -50msec prior to the TMS pulse. Large muscle artefacts, faulty channels, and otherwise bad epochs were removed by visual inspection of the concatenated EEG trace (<10% of total number of epochs selected for rejection). Two rounds of Independent Component Analysis (ICA; FastICA algorithm, ‘tanh’ contrast function and semi-automated component) were implemented via the TESA toolbox in EEGLAB (Delorme & Makeig, 2004; Rogasch et al., 2017). The EEG signal was broken down into separate components via ICA. TESA then categorised each component as consisting of either neural or non-neural activity. The first round of ICA was implemented to identify and remove the large TMS pulse artefact and eye blinks. Once removed, data were bandpass filtered between 1 to 100Hz (4^th^ order Butterworth filter) and 50Hz line noise was removed using a 50Hz notch filter (band-stop filter between 48-52Hz, 4^th^ order Butterworth filter). Data were again visually inspected and bad epochs and channels were removed. A second round of ICA was performed using TESA’s semi-automated selection of neural and non-neural (eye blink/movement and muscle artefacts) components with visual inspection for conformation and rejection of artifactual activity. Missing channels were then interpolated using a spherical interpolation method (EEGLAB; pop_interp function). All channels were re-referenced to an average of all channels. All epochs were then collapsed into a single TEP by averaging each channel over all epochs for every individual and separately for both conditions (L-DLPFC and R-DLPFC). For further information on the TMS-EEG data analysis pipeline used in the current, please see: (Rogasch et al., 2017). An example TMS-EEG analysis pipeline similar to the present study is provided at: https://nigelrogasch.gitbooks.io/tesa-user-manual/content/example_pipelines.html (Rogasch, 2017).

Average TEPs were generated for each subject and separately for each stimulated region of interest (L-DLPFC and R-DLPFC). Analysis focused on the *a priori* determined peaks of interest, namely P60 and N100, which have been suggested to measure excitatory and inhibitory neurotransmission, respectively (Rogasch et al., 2017). To measure the P60 and N100 components the average amplitude over pre-specified time windows was used (P60: 55-75ms; N100: 90-130ms) using automated scripts in Matlab. An analysis was also performed on the ratio of the P60 to the N100. Due to variable voltage shifts across the TEP epoch, some P60 values were negative when measured as averaged amplitude within the window of interest, a feature of the data which would result in non-sensible comparisons of the P60/N100 ratio if not addressed. To address this issue, we visually inspected the data to mark the peak P60 and N100 latencies. We then computed the difference between the peak P60 and N100 amplitude and the average of the minimum of the two surrounding opposite deflections that were within 50ms of the TEP of interest. This provided positive values for all P60 deflections, and negative values for all N100 deflections, representing the size of the deflection compared to the surrounding ongoing TEP.

### Statistical Analysis

To confirm adequate matching between control and meditator groups, the following statistical analyses were performed in SPSS (version 23): Independent samples t-tests were run to test for demographic group matching (age, years of formal education) and matching for scores on self-report measures (BAI, BDI and FFMQ). Chi-square tests were used to verify that groups did not differ in gender and handedness. Prior to statistical analyses Shapiro-Wilk tests and visual inspection of the histograms were conducted to test for normality.

TMS-EEG measures were inspected for outliers using a univariate outlier detection method (data were z-score transformed, and z-scores greater than 3.29 were considered outliers). No outliers were detected using this method for the P60 or N100 amplitude, but one outlier was detected within the control group for the P60/N100 ratio, and this value was winsorized to match the next highest value. To test whether the amplitudes of the P60 and N100 differed between groups, or whether group interacted with hemisphere (i.e., the site of stimulation), two separate mixed model ANOVAs were run in SPSS (version 23). The main effect of condition was not analysed as it was expected from previous research that the left and right sites of stimulation would differ (Farzan et al., 2016), but this provided no valuable insight into the primary measure of interest, i.e. the neural effects of mindfulness meditation. Further analyses of the ratio of the P60 to the N100 (calculated by dividing each individual’s peak P60 deflection by their corresponding peak N100 deflection; 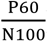) were performed using non-parametric Mann–Whitney U test statistics (Mann & Whitney, 1947). To control for multiple comparisons, the Benjamini-Hochberg ‘False Discovery Rate’ (FDR) method was used (Benjamini & Hochberg, 1995), and p-values adjusted for the FDR are reported as *p*_FDR_. Where null results were obtained, these were complimented by Bayes Factor (BF) analyses to quantify the probability of the null result. BF analyses determines the odds of observing the model describing the *null hypothesis* over that of the model describing the *alternative hypothesis* (expressed as an odds ratio with Bayes Factors > 1 favouring the null hypothesis model; (Rouder et al., 2017)).

To compliment the use of multivariate single electrode and time window analysis, the current study also performed statistical analysis using the ‘Randomization Graphical User Interface’ software (RAGU; (Koenig et al., 2011)). The benefit of using RAGU to analyse EEG data is that it ameliorates several methodological and biasing issues by minimising the need for *a priori* selection of parameters (e.g., electrodes and time windows to analyse). In particular, RAGU uses non-parametric randomised permutation statistical techniques and whole EEG data (all channels, all epochs and all time points) to compare scalp field differences between groups at each time point along the averaged epoch. RAGU employs methods to control for multiple comparisons (Habermann et al., 2018; Koenig et al., 2011) by randomly permuting the data to generate distributions for which the null-hypothesis of no effect holds for each time point. The cumulative size of effects in the real data can then be compared to this distribution to obtain a ‘*global count’ p*-value, which is the probability of observing the real data at all timepoints in the epoch under the assumption that the null-hypothesis is true. This same process can also be applied to obtain a distribution of lengths of contiguous significant time periods in the randomised data termed ‘*duration control*’. Time periods of contiguous significance in the real data that exceed the 95% threshold in the distribution of the randomised data are then highlighted as effects with statistically significant durations in the real data. This duration control ensures the overall level of significance is maintained at the *a priori* determined level of significance (α < 0.05).

To conduct the statistical tests, RAGU uses the Global Field Power (GFP) measure as a reference free measure of global neural response strength. GFP is equivalent to calculating the standard deviation from the average reference over all electrodes. The first step in the RAGU pipeline is to confirm that there exists a consistent activation of neural sources in response to TMS using the Topographical Consistency Test (TCT). This test ensures that significant results in subsequent RAGU analyses cannot be explained by extreme variation within a single group. To determine whether there is topographic consistency, the averaged GFP across all participants within groups and for each time point is calculated (i.e., GFP of the grand-mean map). Values of the grand-mean GFP map depend on both the amplitude and spatial consistency of active neural sources generating the scalp field. If the response to the stimulus is random across participants (i.e., no consistency; random noise) then the grand-mean GFP will average to 0 (or close to 0). However, if there is a consistency in response across individuals, the grand-mean GFP will sum to greater than 0. To statistically analyse these values, they are compared against the null-hypothesis distribution of no consistent effect. This distribution is generated by randomly shuffling the channels within individual observations so that the GFP and variation within individuals remains the same, but when averaged across subjects the grand-mean GFP is now representative of the null-distribution of no consistency (i.e. GFP values for which there is no consistency among the groups). This is then permuted 5000 times. Finally, the grand-mean GFP in the real data is compared to the grand-mean GFP values in the null distribution for each time point, with the proportion of the randomised GFP values exceeding the real observed data forming the *p*-value (the probability of cases in the null distribution taking values larger than the observed grand-mean GFP). The time course of *p*-values then forms the overall TCT indicating periods of time for which the consistency of active neural sources exceeds that of the distribution of no consistency.

Once there is confirmation of a consistent neural activation within groups and conditions using the TCT, further analysis is provided by two separate but related significance tests. The GFP test compared differences in overall neural response strength between groups (meditators and controls) and hemispheres (L-DLPFC and R-DLPFC stimulation) using the GFP as the measure of interest. The between group/condition difference in the GFP of the real data is then compared to the null-distribution of no effect; this distribution is generated by randomised shuffling of the individual observations 5000 times for each measured time point and drawing a distribution of the randomised GFP values. For each time point the real data are compared to the randomised distribution and a *p-*value is determined. Values that exceed the 0.05 alpha-level threshold are considered significant, and the proportion of time points of the real data that exceed this threshold across the whole epoch determines the overall significance (i.e., the global count *p*-value). Contiguous periods of significance across the epoch are then controlled for using global duration statistics.

The GFP test measures differences between groups/conditions of neural response strength. To determine if the topographic distributions of active neural sources differ between groups the Topographical Analysis of Variance (TANOVA) test is performed. Importantly, this test does not require active sources to be localised to cortical regions to determine whether the topographical distributions differ. To achieve this, the TANOVA first applied the L2 normalisation of the data within groups/conditions by dividing the average group/condition voltage values by its scaling factor (the GFP). Any differences between groups/conditions can then be said to be independent of the global strength of neural activity. The within group electrode space data is then averaged, and the average electrode space data from one condition/group is subtracted from the average electrode space data from another condition/group. The *dGFP* is then calculated from this between group/condition normalised difference data. If there are true between-group differences in active brain sources generating the scalp field, then values for the *dGFP* will be large. Statistical analyses are provided by 5000 randomised permutations and the generation of a *dGFP* null-distribution of no effect. The global count *p*-value and duration control statistics are then derived in the same manner as the GFP test.

## Results

### Between group comparisons of demographics and scales

Most demographic variables were found to be normally distributed using Shapiro-Wilk’s test for normality and visual inspection of the histograms. However, scores for the BDI-II and BAI violated tests for normality (**BDI-II**: *Controls*: *W*(19) = 0.836, *p* = 0.004, skewness = 0.854 (*SE =* 0.524), kurtosis = -0.649 (*SE* = 1.014); *Meditators*: *W*(15) = 0.259, *p* = 0.006, skewness = 0.976 (*SE* = 0.580), kurtosis = - 0.096 (*SE* = 1.121); **BAI**: *Meditators*: *W*(15) = 0.832, *p* = 0.010 skewness = 1.347 (*SE* = 0.580), kurtosis = -0.096 (*SE* = 1.121)). Attempts at log transformation (base 10, and the natural log) failed to appropriately normalise the distribution. To mitigate the effect of these violations, comparisons between this data were conducted using non-parametric Mann-Whitney *U* tests, and they were found to be non-significant (**BDI-II**: *U* = 110, *p* = 0.271; **BAI**: *U* = 128, *p* = 0.817). No significant between-group differences were observed in age, gender, years of education or handedness using independent samples t-tests (all *p*-values ≥ 0.097; results provided in Table 1). A statistically significant difference was, however, observed between groups on the FFMQ, with experienced meditators scoring an average 24.49 points higher compared to controls (*t*(32) = -4.130, *p* < 0.001, 95% CI [-36.57, -12.41], *d* = 1.452). Demographic measures and results are presented in Table 1.

**Table 1.** Reports of Demographic Variables, Including Measures of Central Tendency and Statistics. *Note:* SD = Standard Deviation; FFMQ = Five Facet Mindfulness Questionnaire; BAI = Beck Anxiety Inventory; BDI = Beck Depression Inventory, *χ*^2^ = chi-square test statistic, t = independent samples t-test statistic, U = Mann-Whitney U statistic for non-parametric tests. All significance levels (α) set at 0.05.

|  | Controls<br>*M (SD)* | Meditators<br>*M (SD)* | Statistics |
| --- | --- | --- | --- |
| Age | 29.58 (11.48) | 35.67 (13.81) | $t(32) = 1.404, p = 0.170$ |
| Gender (Male/Female) | 10 / 9 | 12 / 3 | $\chi^2(1) = 2.749, p = 0.097$ |
| Years of Education | 17.31 (2.04) | 17.07 (2.71) | $t(32) = 0.338, p = 0.737$ |
| Handedness<br>(Right/Left/Mixed) | 15 / 3 / 1 | 15 / 0 / 0 | $\chi^2(2) = 3.579, p = 0.167$ |
| Years Meditating | - | 6.23 (4.55) | - |
| Hours Spent<br>Meditating Per Week | - | 7.49 (7.00) | - |
| FFMQ | 132.84 (19.12) | 157.33<br>(14.26) | $t(32) = 4.130, p < 0.001$ |
| Independent Samples Non-Parametric Mann-Whitney U Tests |  |  |  |
| BAI | 5.06 (3.83) | 5.80 (6.54) | $U = 128.000, p = 0.817$ |
| BDI-II | 4.32 (4.66) | 2.40 (2.95) | $U = 110.000, p = 0.271$ |

### Single Electrode Analyses

#### TMS-Evoked Potentials

The group-averaged single-electrode TEPs are provided in Figure 1. Butterfly plots of the group averaged TEPs are depicted in Figure 2, revealing characteristic TEP deflections at N45, P60, N100, P200 and N280. The TEPs displayed classic waveforms in response to DLPFC stimulation (Lioumis et al., 2009).

**Fig. 1.**
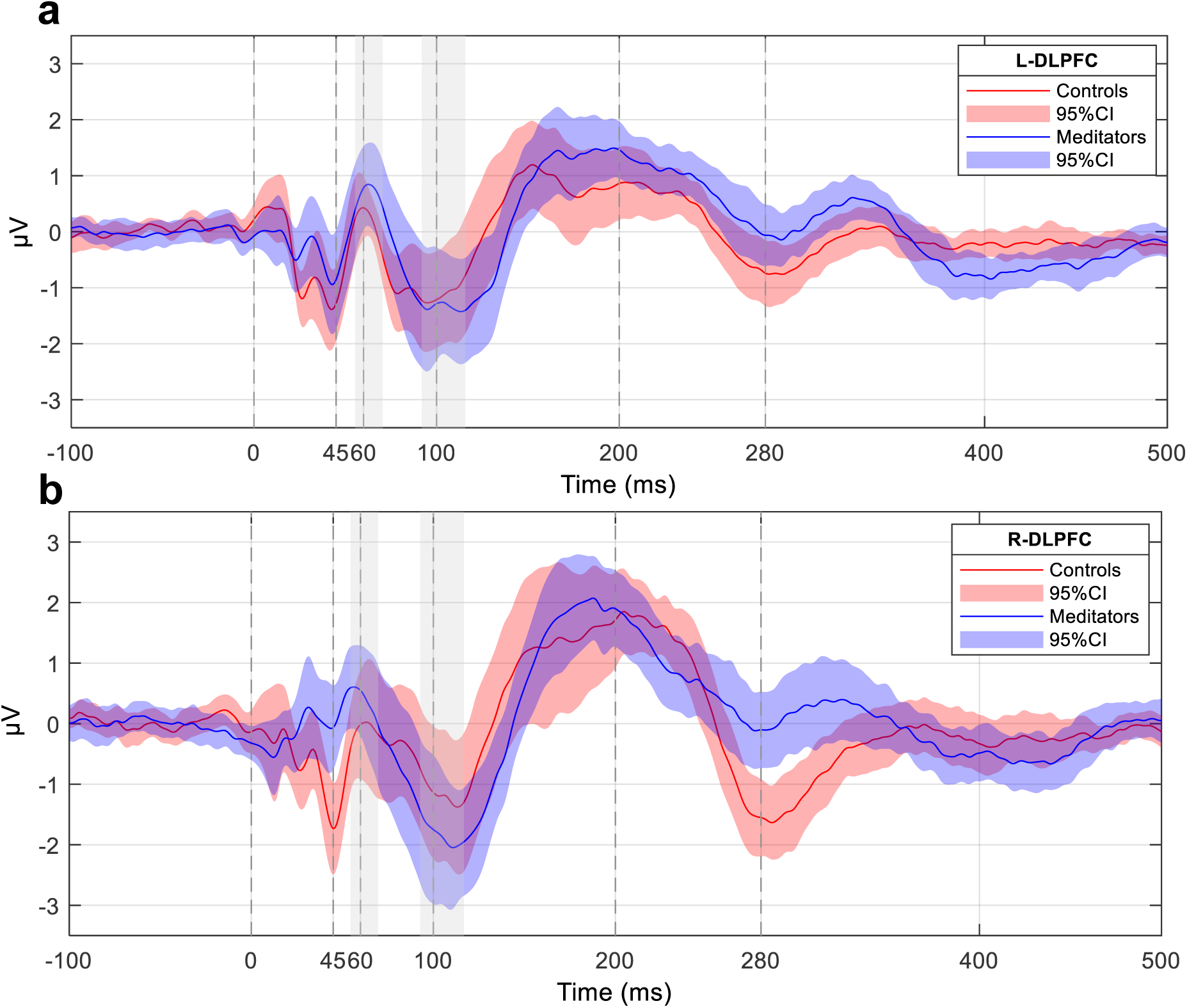
Group Averaged TMS-Evoked Potentials from Single Electrodes: (**a**: F3; Left-DLPFC) and (**b**: F4; Right-DLPFC) show the potentials evoked post-TMS for meditators (n = 15, blue waveform) and controls (n = 19, pink waveform). Transparent shading signifies the 95% Confidence Interval (CI). Shaded grey sections mark the epochs utilised to compute the averaged peak amplitude for P60 and N100 components. The group TEPs represent amplitude (µV) changes from baseline across time (ms), with TMS administered at time = 0. Black dashed lines denote standard latencies for respective components (N45, P60, N100, P200, and N280)

**Fig. 2.**
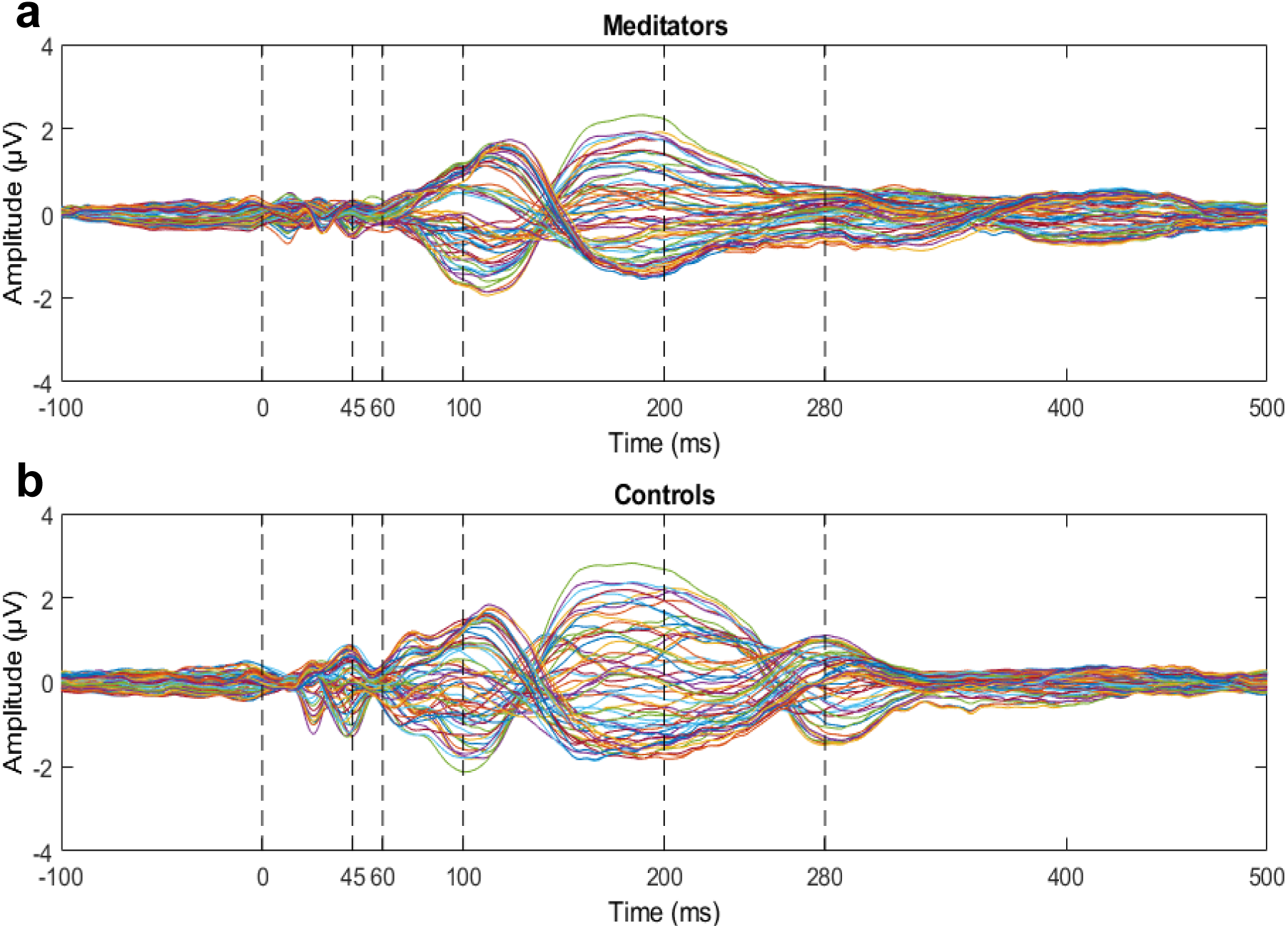
Group-Averaged Butterfly Plots of TMS-Evoked Potentials (TEPs): These plots showcase TEPs for both left and right dorsolateral prefrontal stimulation in meditators (**a**, n = 15) and controls (**b**, n = 19). TEPs are represented as amplitude changes (µV) from baseline across time (ms), with TMS applied at time = 0. Dashed lines denote the standard TEP components: N45, P60, N100, P200, and N280

To investigate whether P60 and N100 differed between groups, mixed model ANOVAs were employed using data obtained from the single electrodes of the stimulated sites (F3 and F4). Preliminary analysis of the separate mixed model ANOVAs of P60 and N100 amplitudes indicated that most measures of normality and homogeneity of variance were not violated (*F_max_*, Box’s *M,* Mauchly’s Test of Sphericity). However, the assumption of between group equality of error variance in P60 amplitudes was violated (Levene’s test for equality of error variances: L-DLPFC, *F*(1, 32) = 0.014, *p* < 0.05; R-DLPFC *F*(1, 32) = 0.231, *p* < 0.05). Unequal group sizes, as in the present study, further exacerbate deviations away from between group equality of variance. Despite this violation, the mixed model ANOVA *F*-test has been shown to be robust against deviations from unequal group sizes if the disparity between them are small (largest group size/smallest group size < 1.5; current study 19/15 = 1.27; (Stevens, 2012)). Corrections for violations of the equality of error variances are limited with respect to mixed model ANOVAs. Applying a lower alpha threshold (α < 0.05) is one suggested method to control for the inflation of a type-I error rate (Allen & Bennett, 2007). However, the current (and subsequent) ANOVA analyses show that no group or factor measures met the requirements for statistical significance, and methods for correcting for this would favour the results towards the null hypothesis of no effect. The interpretations of these null results without correction are therefore warranted.

No significant main effect of group in P60 amplitudes were observed (*Group; F*(1, 32) = 1.712, *p* = 0.200, η^2^ = 0.051). In addition, no significant interaction effect between group and hemisphere was observed (*Group*Hemisphere; F*(1, 32) = 0.303, *p* = 0.586, η^2^ = 0.003). No significant differences in N100 amplitudes were observed between groups (*Group; F*(1, 32) = 1.128, *p* = 0.296, η^2^ = 0.034). In addition, no significant interaction effect between group and hemisphere was observed (*Group*Hemisphere*; *F*(1, 32) = 0.130, *p* = 0.721, η^2^ = 0.001). The results of the ANOVA tests are displayed in Table 2.

**Table 2.** Mixed-Model ANOVA Results of the Single Electrode Comparisons for Between Groups and Sites of Stimulation. *Note*: M = Mean; SD = Standard Deviation; FDR = False Discovery Rate (displayed with the adjusted alpha level α_adj_); η_p_^2^ = partial eta squared, η^2^ = eta squared; BF_01_ = Bayes Factor Analysis of the Null Hypothesis (BF_01_ > 1 favours the model of the null hypothesis). All significance alpha levels set at 0.05.

| Condition | Controls<br>*M* (*SD*) | Meditators<br>*M* (*SD*) | Statistics |
| --- | --- | --- | --- |
| P60 L-DLPFC<br>(F3 electrode) | -0.118<br>(0.79) | 0.580<br>(1.48) | **Main effect of group:** $F(1, 32) = 1.712, p = 0.200$ ($p_{FDR} = 0.504$), $\eta_p^2 = 0.051$, $\eta^2 = 0.051$, $BF_{01} = 1.768$<br>**Main effect of hemisphere:** $F(1, 32) = 0.303, p = 0.586$ ($p_{FDR} = 0.704$), $\eta_p^2 = 0.009$, $\eta^2 = 0.004$, $BF_{01} = 3.620$ |
| P60 R-DLPFC<br>(F4 electrode) | -0.133<br>(2.01) | 0.228<br>(1.53) | **Interaction:** $F(1, 32) = 0.256, p = 0.616$ ($p_{FDR} = 0.704$), $\eta_p^2 = 0.008$, $\eta^2 = 0.003$, $BF_{01} = 6.306$ |
| N100 L-DLPFC (F3<br>electrode) | -0.791<br>(2.04) | -1.346<br>(1.88) | **Main effect of group:** $F(1, 32) = 1.128, p = 0.296$ ($p_{FDR} = 0.504$), $\eta_p^2 = 0.034$, $\eta^2 = 0.034$, $BF_{01} = 1.591$<br>**Main effect of hemisphere:** $F(1, 32) = 1.044, p = 0.315$ ($p_{FDR} = 0.504$), $\eta_p^2 = 0.032$, $\eta^2 = 0.006$, $BF_{01} = 2.714$ |
| N100 R-DLPFC (F4<br>electrode) | -0.983<br>(2.18) | -1.747<br>(1.74) | **Interaction:** $F(1, 32) = 0.130, p = 0.721$ ($p_{FDR} = 0.721$), $\eta_p^2 = 0.004$, $\eta^2 = 0.001$, $BF_{01} = 4.219$ |

The P60/N100 ratios violated the required assumption of normality in parametric repeated measures ANOVAs using the Shapiro-Wilk test (all *p* < 0.017). To control for the violation of these assumptions, Mann–Whitney U test statistics (Mann & Whitney, 1947) were used to test for differences in the P60/N100 ratios (results presented in Table 3). Significant between group differences were observed in the L-DLPFC condition (U = 270.5, *p* < 0.001, *d* = 1.571, BF_10_ = 50.67, significant at FDR adjusted *p*_FDR_ *= 0*.004) and R-DLPFC condition (U = 252.5, *p* < 0.001, *d* = 1.447, BF_10_ = 39.24, significant at FDR adjusted *p*_FDR_ *= 0.004*). Meditators were observed to have higher P60/N100 ratios on average compared to controls, reflecting that their P60 deflections were on average more similar in size to their N100 deflections compared to the control participants (who showed N100 deflections that were larger than their P60 deflections, see Figure 3). Finally, to confirm differences in the P60/N100 ratios were not simply the result of differences in either the P60 or N100 when these TEPs were measured as deflections (rather than as averaged amplitudes within the window of interest as per our amplitude analyses), we performed two confirmatory repeated measures ANOVAs comparing the P60 and N100 deflection values that were used in computing the P60/N100 ratios. These analyses showed no significant differences between the groups or interactions between group and hemisphere in the P60 or N100 deflections (all *p* > 0.18). This result aligns with our null result for the P60 and N100 amplitudes when measured averaged across their windows of interest and indicates that the P60/N100 ratio effect is driven by between group differences in the relationship between the P60 and N100 ratio.

**Fig. 3.**
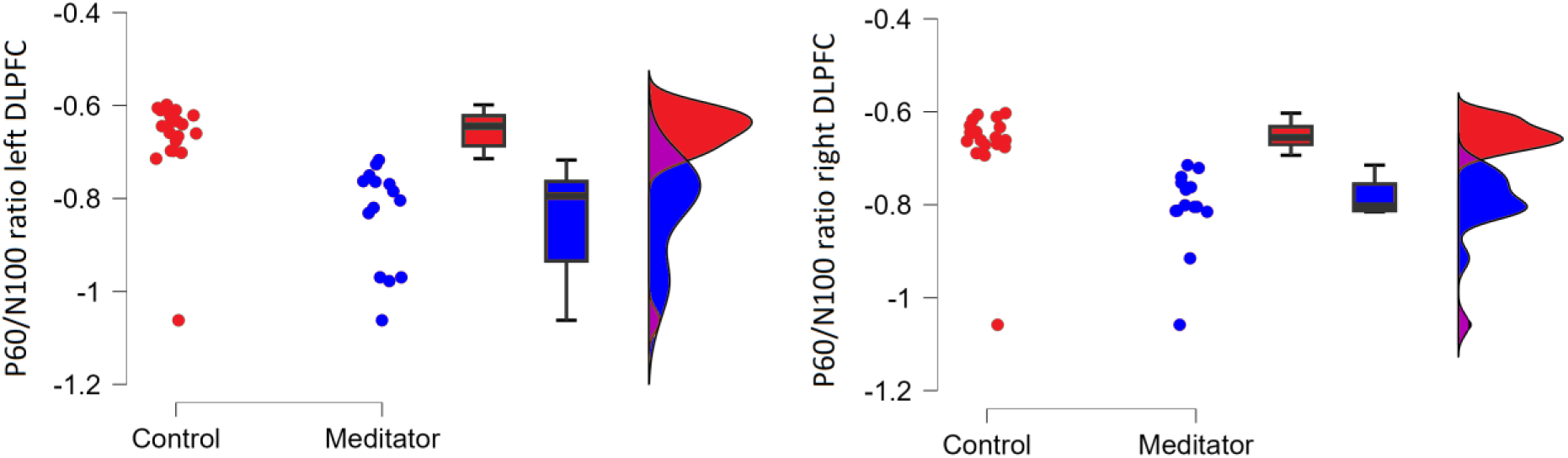
Raincloud plots of the peak P60/N100 ratios for the left and right dorsolateral prefrontal cortex (DLPFC). Note that all N100 peak values were negative, resulting in negative values for all P60/N100 ratios. Significant between group differences were observed for both the left (U = 270.5, p < 0.001*, d = 1.571 [p_FDR_ = 0.004], BF_10_ = 50.67) and right DLPFC (U = 252.5, p < 0.001*, d = 1.447 [p_FDR_ = 0.004], BF_10_ = 39.24). Raw P60 and N100 deflection values used in calculating the P60/N100 ratios can be found in Supplementary Materials, Figure 1

**Table 3.**
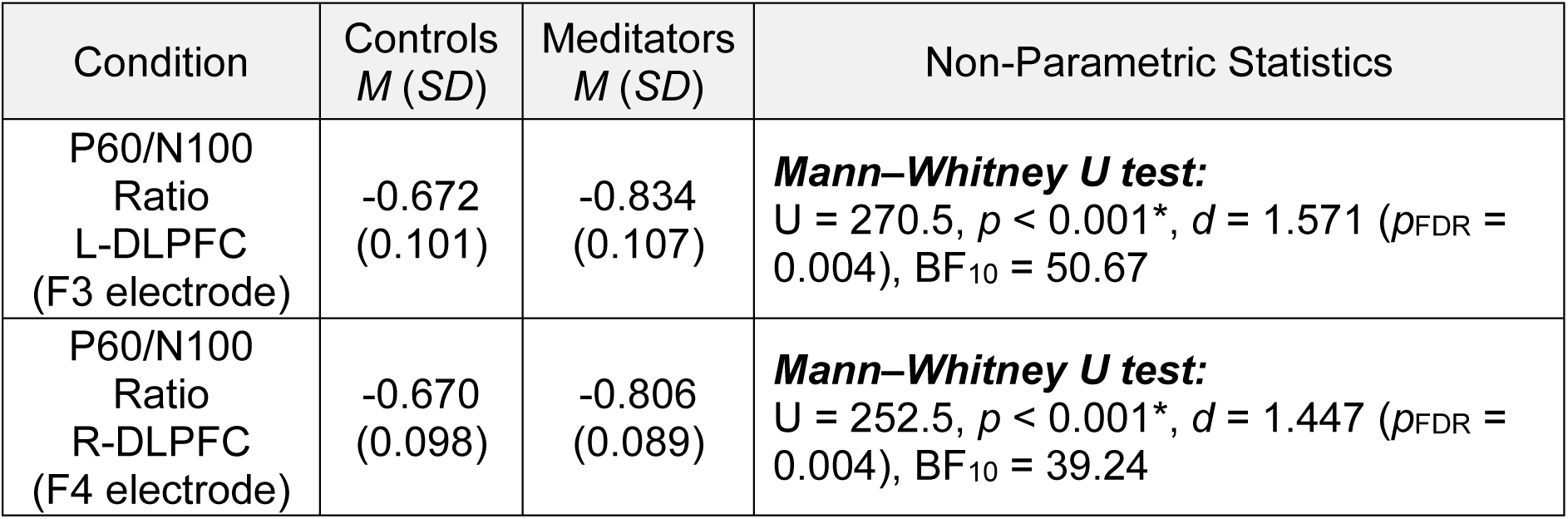
Mann–Whitney U Test Results for Between Group Analyses of the P60/N100 Ratios Using Non-Parametric Analyses. *Note:* M = Mean; SD = Standard Deviation; FDR = False Discovery Rate (displayed with the adjusted alpha level α_adj_), d = Cohen’s measure of effect size determined using an independent samples t-test. BF_10_ = Bayes Factor Analysis of the Alternative Hypothesis (BF_10_ > 1 favours the model of the alternative hypothesis). All alpha significance levels set at 0.05. Note that the tests remain significant after FDR correction. *reflects p-values that were significant at p < 0.001.

| Condition | Controls<br>*M (SD)* | Meditators<br>*M (SD)* | Non-Parametric Statistics |
| --- | --- | --- | --- |
| P60/N100 Ratio<br>L-DLPFC<br>(F3 electrode) | -0.672<br>(0.101) | -0.834<br>(0.107) | ***Mann–Whitney U test:***<br>$U = 270.5$, $p < 0.001^*$, $d = 1.571$ ($p_{FDR} = 0.004$), $BF_{10} = 50.67$ |
| P60/N100 Ratio<br>R-DLPFC<br>(F4 electrode) | -0.670<br>(0.098) | -0.806<br>(0.089) | ***Mann–Whitney U test:***<br>$U = 252.5$, $p < 0.001^*$, $d = 1.447$ ($p_{FDR} = 0.004$), $BF_{10} = 39.24$ |

### Whole Scalp Field Analysis of TMS-evoked potentials

The TCT test showed that there was consistency in the active neural sources contributing to the scalp field throughout most of the epoch (-100 to 500ms) for both groups and conditions (Figure 4; overall *p* = 0.0002). There were, however, short periods of inconsistent neural activity that were observed in all groups and conditions. Inconsistent activity can be observed in periods within the P60 time window used in the single electrode analysis (55-75ms). Inconsistencies were also observed for meditators in the R-DLPFC condition between 280-287ms, and for controls in the L-DLPFC condition between 312-347ms. Inconsistencies in these periods indicate that significant results obtained in the TANOVA test (described in subsequent analyses) may be due to inconsistency in the topographical distribution of the neural response across individuals within a single group.

**Fig. 4.**
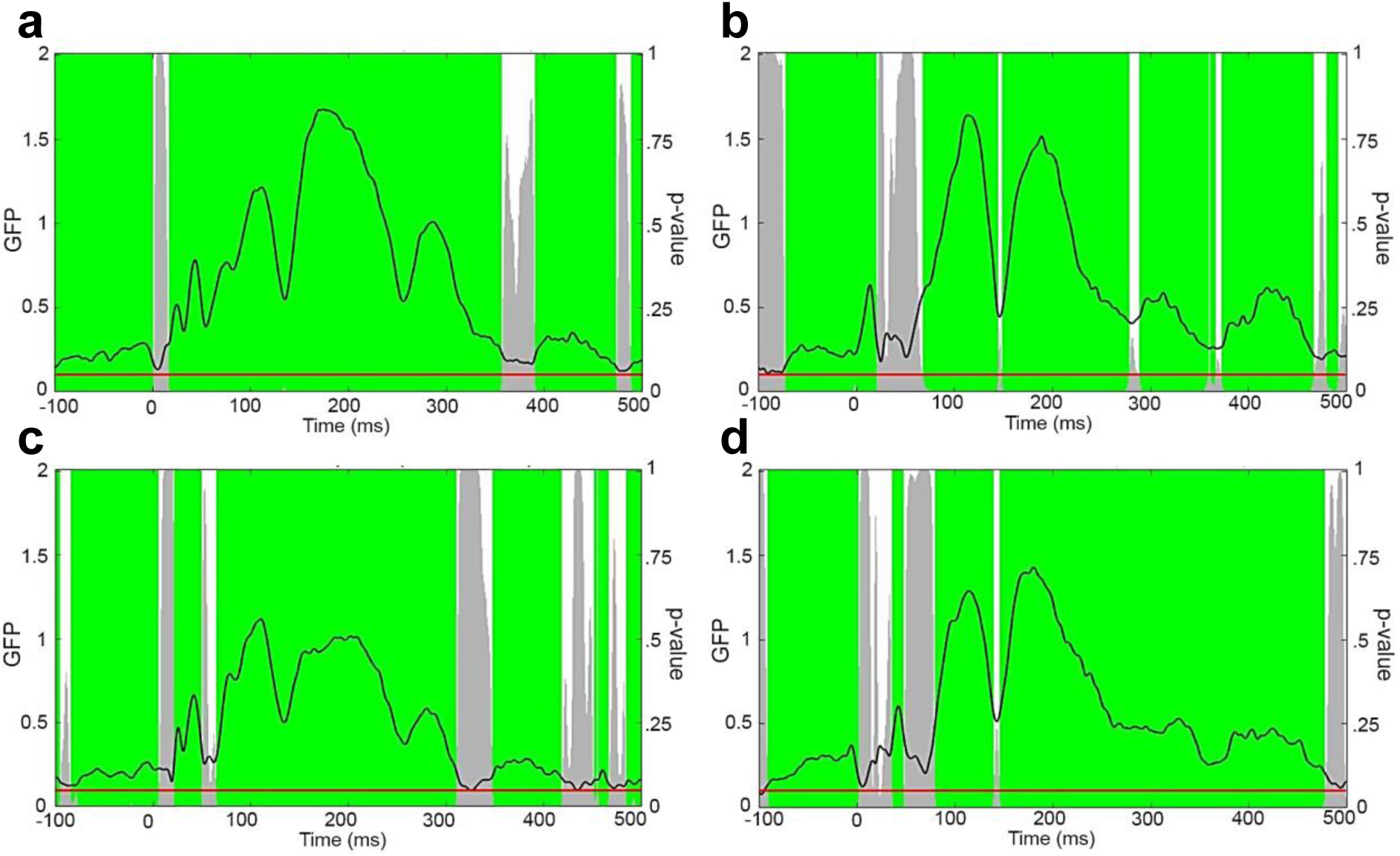
Topographical Consistency Test of Neural Response to TMS: Comparisons between groups and conditions are presented: **a** and **c** for controls; **b** and **d** for meditators. Conditions correspond to **c** and **d** for left dorsolateral prefrontal cortex post-stimulation at the F3 electrode, and **a** and **b** for right dorsolateral prefrontal cortex post-stimulation at the F4 electrode. Graphs display Global Field Power (GFP) variations over time, with TMS delivery marked at time = 0. The GFP measure, involving the rectification of the TEP, results in the distinct waveform appearance shown. The p-threshold is indicated by a solid red line (0.05). White sections denote non-significant periods (p < 0.05) with inconsistent neural source activations, while grey contours reflect the p-value’s temporal trajectory. Green regions highlight consistent neural activity, predominantly observed between 80-310ms. Inconsistencies are evident for meditators from 0-80ms and controls between 310-390ms

The between-group GFP tests for differences in neural response strength were non- significant (group main effect, global count *p* = 0.485, see Figure 5). Additionally, no significant interaction between group and condition was observed (hemisphere*group interaction, global count *p* = 0.610). Further, the brief periods of significance did not survive multiple comparison duration control (global duration statistics: main effect of group = 39 ms; hemisphere*group interaction = 30 ms). Of note, no periods of significance were observed around the time windows of interest in the hypothesis driven analyses of P60 and N100 in agreement with the single electrode analyses. Results of the GFP test indicate that there were no differences between groups in the overall strength of neural response to TMS to the DLPFC for either hemisphere.

**Fig. 5.**
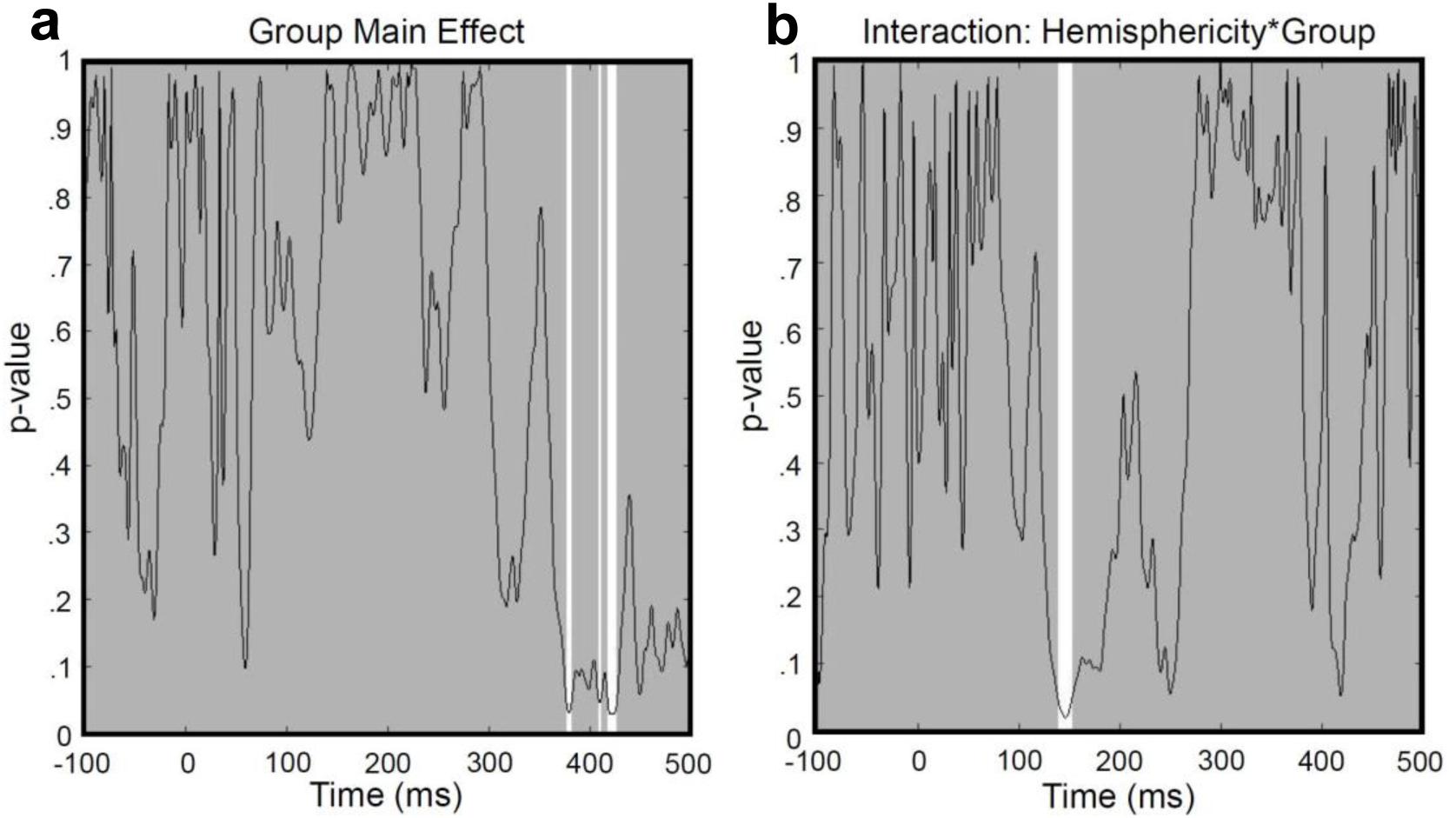
Comparative Analysis of Neural Response Strength to Single-Pulse TMS: Controls vs. Meditators: Depicted above are the results of the (**a**) between-group statistical comparisons of Global Field Power (GFP) between controls and meditators, and (**b**) the interaction effects based on the site of TMS stimulation, focusing on left and right dorsolateral prefrontal cortices. The x-axis maps time, with TMS delivery marked at time = 0. The evolving p-value is traced by the black line across the timeline. Grey sections represent periods where the observed difference in GFP (dGFP) aligns within the 95% non-significant range of the randomised null-hypothesis distribution. In contrast, white sections highlight periods where dGFP values exceed the critical significance threshold (p < 0.05), yet fall short of the duration criteria for multiple comparison significance. Notably, no significant variations in neural response strength were detected in the analyses

In contrast to the null results in the GFP test, significant group differences in the distribution of active neural sources were observed in the TANOVA test lasting from 269 to 332ms post TMS pulse (global count *p* = 0.016; Figure 6a). This period was the only period to survive the multiple comparisons duration control of 32ms (the main effect of group averaged significance and effect size between 269-332ms were: *p* = 0.008, η^2^ = 0.126). No significant interaction between group and condition survived the multiple comparisons duration control of 25ms (hemisphere*group interaction, global count *p* = 0.655). The significant group effect between 269-332ms post TMS pulse is evidence that there were different spatial distributions of active neural sources contributing to the scalp field during this time. However, as topographical inconsistencies were observed in the TCT (meditators between 280- 287ms; controls between 312-347ms), at least some of this effect may have been due to within group variation. Although there was no interaction effect, the TCT showed a different pattern of topographical consistency for stimulation to the different hemispheres. As such, we examined whether the significant main effect of group replicated within a single hemisphere that showed consistent topographical activation. The results of this analysis are provided in Figure 6c and 6d. No between-group significance was present in the L-DLPFC condition (global count *p* = 0.289), and no significant periods survived the multiple comparisons duration control of 31ms (Figure 6c). A significant between-group difference was observed within the R- DLPFC stimulation condition (global count *p* = 0.029) which lasted between 270- 332ms post TMS (η^2^ = 0.097) and survived the multiple comparisons duration control of 31ms. Inconsistencies observed in the TCT with R-DLPFC stimulation were only observed in meditators between 280-287ms (Figure 6d). The remaining time window (287-332ms) still exceeded the multiple comparisons duration control of 31ms. This result suggests that the meditation group showed a different topographical distribution of active neural sources following R-DLPFC stimulation. Further characterisation of the topographical distribution of neural activity over the period of significance is provided in Figure 6.

**Fig. 6.**
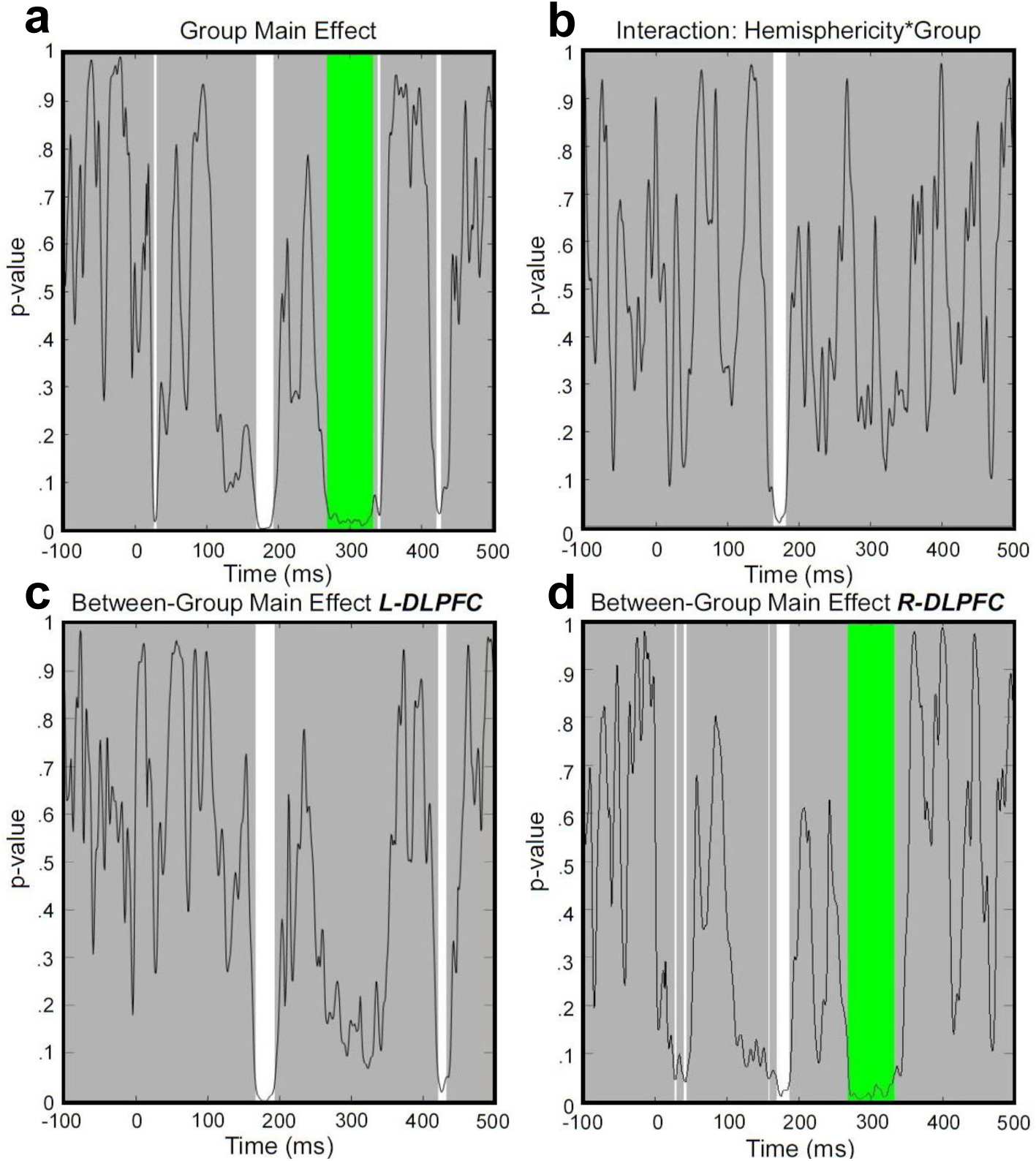
Topographical Analysis of Variance (TANOVA) for Neural Activity Distribution: This figure illustrates (**a**) between-group comparisons of controls versus meditators, (**b**) interactions based on the stimulation site (L-DLPFC vs. R-DLPFC), (**c**) isolated between-group comparisons for L-DLPFC stimulation, and (**d**) isolated between-group comparisons for R-DLPFC stimulation. The timeline on the x-axis starts with TMS delivery at time = 0, whilst the evolving p-value is marked by the black line on the y-axis. Grey sections denote periods with no statistically significant effects. White sections signify periods of distinct neural source contributions to the scalp’s electromagnetic field without meeting the significance duration threshold. Notably, green highlights pinpoint periods meeting the 32ms duration threshold, with a significant group variance observed between 269-332ms post-stimulation in section **a**

The topographical distributions of neural activity averaged across 269 to 332ms post TMS pulse showed less fronto-midline negativity and greater posterior negativity in meditators (Figure 7a) when compared to controls (Figure 7b). The meditator group also showed greater central and temporal positivity compared to controls. In the control group, the positive voltages appeared to be shifted more posteriorly, focused over centro-parietal regions. In addition, controls appeared to show more equivalent centro-parietal positivity across both hemispheres compared to meditators (who exhibited greater positive voltages in electrodes over the left hemisphere). The t-map (Figure 7c) illustrates the difference (controls subtracted from meditators) in voltage distributions during the significant period. Specifically, the t-map indicates that the differences between groups appear to be strongest around the general region of the left-sided site of stimulation (F3, F5 and AF3 electrodes) with meditators showing more positive neural activity compared to controls. In addition, meditators showed more negative voltages compared to controls in parieto-occipital regions (PO3 and Pz electrodes).

**Fig. 7.**
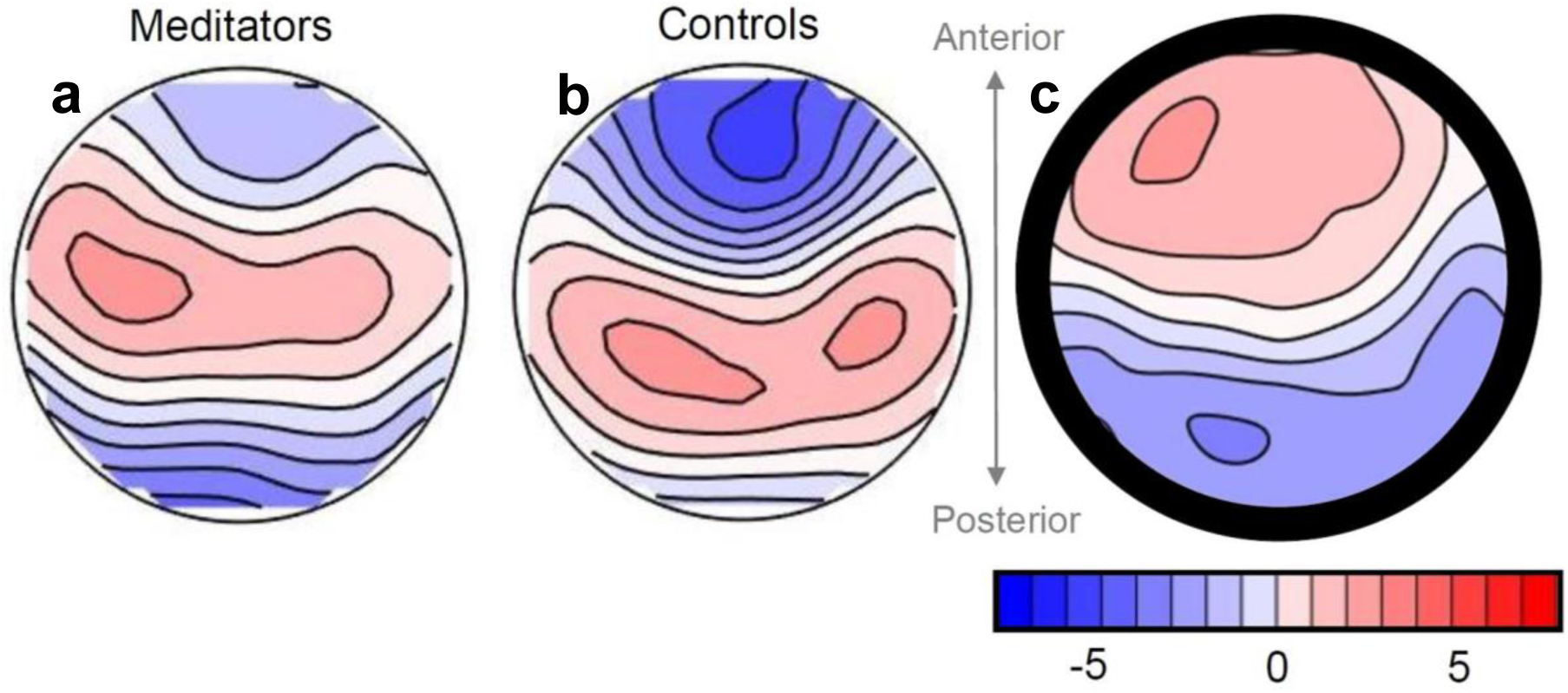
Between-group Averaged Topographies during Significant TANOVA Period (269 – 332ms post-stimulation, p = 0.008, η^2 = 0.126): Displayed are group- averaged topographies for (**a**) meditators and (**b**) controls. The differential topographical map (**c**, t-map) is calculated by subtracting the controls from the meditators, illustrating the principal topographic contribution of the meditator group. The colour scale signifies the strength of the time-window averaged polarity for each electrode. Notably, meditators exhibit reduced negative activity in frontal regions. Their positive polarity is more centrally located, and they demonstrate heightened posterior negativity compared to the control group.

## Discussion

The current study aimed to use TMS-EEG measures of cortical reactivity to assess whether the DLPFC of individuals with an average of 6.23 years of practicing meditation shows alterations compared to demographically matched non-meditators. Our results indicated that meditators demonstrated significantly higher P60/N100 ratios compared to matched controls, with strong Bayesian evidence despite the small sample size. Interestingly, comparisons between the P60 and N100 components themselves did not significantly differ. This null result for either TEP in isolation coupled with a larger P60/N100 ratio may indicate that meditation is related to a shift in the balance between the two TEPs, but not an effect that is specific to only one of the TEPs. Additionally, the meditation group showed a difference in the distribution of active neural sources 269-332ms following DLPFC stimulation. Analysis of stimulation of L-DLPFC and R-DLFPC indicated that the period 270- 332ms post TMS remained significantly different between the groups and withstood multiple comparisons duration control with R-DLPFC stimulation. This late TEP activity has been suggested to reflect slow inhibitory post synaptic potentials or longer cortico-subcortical-cortical reverberation in response to TMS (Ferreri et al., 2011).

With regards to the interpretation of our results, the putative difference in the DLPFC’s E/I balance in experienced meditators could be explained with reference to the influence of broader alterations in brain anatomy. In particular, a recently proposed model of the mechanisms underpinning the effects of focused attention mindfulness meditation, suggests that the DLPFC and other executive control regions activate to over-ride default thought processes (Ganesan et al., 2022). Greater connectivity between the DLPFC and regions of the ventral and dorsal- corticolimbic networks (which are involved in top-down cognitive control processes) has been observed following a mindfulness meditation intervention (Taren et al., 2017). This top-down control is proposed to reduce the intensity of internal thought processes and allow focus to be sustained on interoceptive sensations (Ganesan et al., 2022). Recent evidence has highlighted the critical importance of E/I ratio in determining neural network dynamics that either improve or degrade local and interregional information transfer (Ma et al., 2019). As such, our current results may point to an altered E/I ratio in the DLPFC as a potential neural candidate by which meditation can lead to enhancements in top-down control by the DLPFC, which enable increased regulation of other brain regions, reduced mind wandering. and improvements to cognitive processes (e.g., executive attention network functions). We note that previous research has identified a link between increased P60/N100 ratio and working memory performance, providing support for this suggestion (Chung et al., 2019). Recent evidence also suggests that the strongest different in the resting brain activity of meditators is an increased in the temporal stability of the EEG signal in parietal regions (responsible for self-awareness, attention and somatosensory integration), suggesting the potential for an integrated view of meditation related changes to the brain, with increased regulation by the DLPFC causing more stable activity in other regions (Bailey et al., 2023e). Further research is recommended to probe this suggested connectivity model of the effects of meditation on brain activity.

Several clinical disorders, including anxiety and depression, are believed to be associated with E/I ratio uncoupling and the resulting disordered prefrontal oscillatory phase coherence (Lisman, 2012; Radhu et al., 2014; Voineskos et al., 2019; Yizhar et al., 2011). The altered E/I ratio following TMS applied to the DLPFC of meditators may suggest a potential target for therapeutic interventions, where therapies could aim to adjust the E/I ratio in a way that might relate to improvement of clinical symptoms following a mindfulness intervention (Piet & Hougaard, 2011; Strauss et al., 2014), Interestingly, recent research has suggested that a steeper N100 slope (driven by a larger P60 amplitude) is found in individuals who respond to TMS treatment for depression, a finding that may be analogous to our finding of a larger P60/N100 ratio in the current research, highlighting a potential clinical implication of the current study (Bailey et al., 2023d).

In contrast to the effects for the P60/N100 ratio, our no differences were detected in the P60 and N100 peak amplitudes individually. As such, the results of the present study may suggest that rather than a physiological effect on either excitation or inhibition, mindfulness meditation has a physiological effect on the processes which determine the balance between excitation and inhibition. It is worth noting that the generation of individual TEP components are not governed by functionally separate neurophysiological processes (Hill et al., 2016), and processes involved in altering the E/I ratio are also implicated in the generation of individual P60 and N100 amplitudes.

The 269-332ms period of significance following DLPFC stimulation found in the mindful meditator group indicates differences in the neurophysiological processes leading to the formation of the N280 component with a peak latency falling between 266.7 ± 32.2ms (Ferreri et al., 2011) and 308.3 ± 7.8ms (Määttä et al., 2019). Polysynaptic circuits with potentials that are mediated via the generation of slow inhibitory post synaptic potential caused by GABA_B_/NMDA receptor mediated neural transmission have been suggested to result in the long latency and wide cortical distributions (measured using EEG) of the N280 component (Farzan et al., 2013). These potentials may also arise from longer cortico-subcortical-cortical reverberation in response to TMS (Ferreri et al., 2011). Studies showing complete cessation of TEPs at around 150ms post stimulation in unconscious individuals (Ferrarelli et al., 2010; Massimini et al., 2005; Rosanova et al., 2012; Sarasso et al., 2015), providing evidence that the higher-order prefrontal cognitive activity contribute to the N280 potential. However, previous characterisation of this component has primarily been in response to stimulation of M1 (Ferreri et al., 2011). Research has also suggested that a common neural mechanism exists between the generation of N100 and N280 (Farzan et al., 2013) due to correlation between the N100, N280 and the length of the CSP (an indirect marker for GABA_B_ergic neurotransmission). However, previous research has only examined voltage amplitudes at single electrodes or clusters of electrodes with relevance to the N280. As such, it is not clear what the functional interpretation of an altered distribution of activity for the N280 in experienced meditators reflects. Candidate explanations might be altered neural connectivity which alters the cortico-subcortical-cortical reverberatory response to the TMS pulse, altered slow inhibitory processes, or other, as yet uncertain explanations.

As far as we are aware, the current study is the first study to assess TMS-EEG applied to the DLPFC in experienced meditators. Previous research that used TMS- EEEG to examine M1 indicated enhanced GABA_B_ergic inhibitory neurotransmission in meditators (Guglietti et al., 2013). However, this previous research, applied stimulation after a 60 minute meditation practice, so their findings may reflect a state effect rather than a trait effect (Tang et al., 2016). Further, changes at M1 in meditators may reflect alterations in somatosensory processes, for example in self-related interoceptive sensory processing, which is known to be modified by meditation (Tang, 2017), rather than attention and executive function related brain regions which are more likely probed by the DLPFC stimulation in the current study.

## Limitations and Future Directions

The functional interpretation of TEPs is controversial and still debated, with research suggesting that the P60 reflects excitatory synaptic activation and the N100 inhibitory processes, while other research suggests the TEPs are also influenced by auditory processing of the TMS click (Conde et al., 2019; Ter Braack et al., 2015). The use of the sound masking protocol in the current study has been shown to reduce the influence of auditory processing on the N100, but residual influence is still suggested to remain (Biabani et al., 2021; Biabani et al., 2019; Conde et al., 2019). While this controversy has implications for the interpretation of our results, it does not obviously invalidate our finding of an altered P60/N100 ratio, as it is not clear why auditory processing related effects would differ between meditators and non-meditators, and why any potential difference would influence the P60/N100 ratio. In particular, our previous research that examined neural activity in response to an auditory oddball using some of the same participants that were included in the current study indicated no differences between the meditator and non-meditator groups in auditory processing ERPs (Payne et al., 2020).

Secondly, variability in the definition of the term ‘mindfulness’ and the inclusion of heterogenous samples of meditators from different traditions are common methodological issues in meditation studies (Van Dam et al., 2018). These factors were controlled for in the current study to some extent by using an inclusion criteria that limited the meditation group to participants who practice breath/body focused meditation (Creswell, 2017; Norris et al., 2018; Tang, 2017; Tang et al., 2015). Additionally, some research has argued that distinctions between different mindfulness meditation practices may not provide much in the way of meaningful differences to neurophysiology, that the differences between meditators and non-meditators is likely to be considerably larger, and that there is conceptual ambiguity in the definition of different meditation types and how these factors might affect neurophysiology (Bailey et al., 2023a-b; Schoenberg & Vago, 2019). Nonetheless, future research could reduce potential heterogeneity from the inclusion of a range of meditation practices by restricting recruitment of meditators to those who practice only a focused attention form of meditation for example.

Another common limitation of meditation research, including the current study, is the use of a cross-sectional study design, and the associated limitation in our ability to attribute causation in the differences associated with the meditator group. This limitation could be addressed by employing a controlled long-term prospective study. However, doing so would be difficult given the effects observed in the present study are associated with an average of 6.23 years of practicing meditation. Having said that, structural and functional brain alterations have been observed after mindfulness interventions ranging from 3 days to 8 weeks (Taren et al., 2017; Tomasino & Fabbro, 2016; Xue et al., 2011).

In addition to the cross-sectional nature of our study, sample selection factors might have affected our results. Although the two groups did not significantly differ in age, the meditator group’s mean age was older. This factor may have increased within group variability of the N100 amplitudes, and it is possible it may have influenced the null result for differences in N100 amplitudes. Future research could address this by directly age matching participants from each group.

Finally, our study had a limited sample size. However, the use of Bayesian analyses indicated that our sample combined with the observed effect sizes provides moderately confidence in our conclusions for many of the null results, and strong confidence in our conclusions for the two positive results. However, future research with a larger sample size will provide more confidence and increase the potential generalisability of results.

## Acknowledgements

We gratefully acknowledge the Dhamma Aloka Vipassana meditation centre in Melbourne and the Melbourne Insight Meditation centre for their assistance with the recruitment of several meditators who took part in the study.

---

**Supplementary Fig 1.**
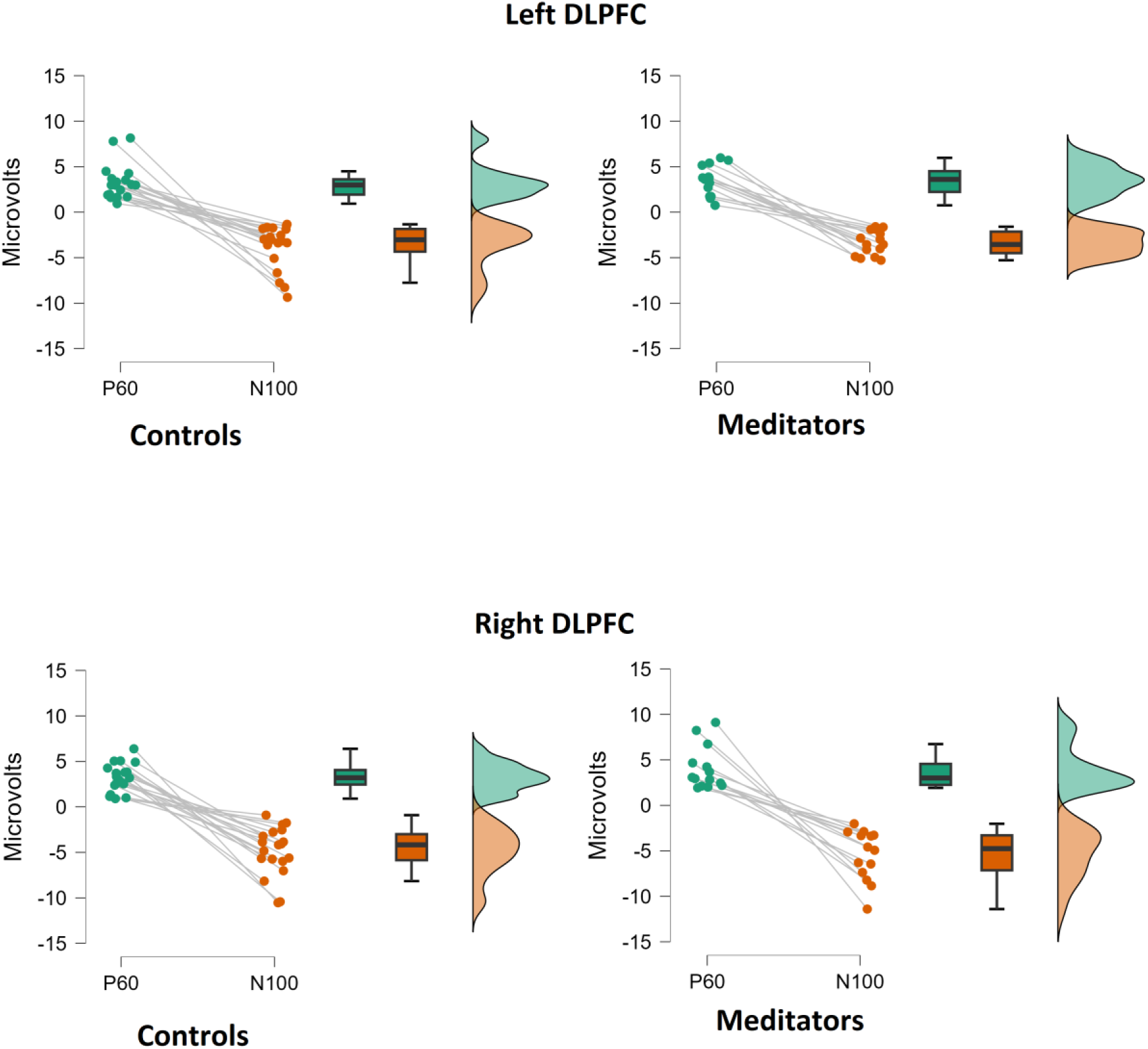
P60 and N100 deflection amplitudes (calculated as the difference between the peak of the P60 and N100 deflection and the mean of the largest opposite value to the peak of the deflection within 50ms of the peak on each side of the peak). These values were used to compute the P60/N100 ratio.

